# Mechanical properties of the unguitractor apodeme of sunny stick insects, *Sungaya aeta* (Phasmatodea: Heteropterygidae)

**DOI:** 10.1101/2025.08.12.669824

**Authors:** Mia Yap, Frederik Püffel, David Labonte

## Abstract

An integral component of musculoskeletal systems are elastic elements mechanically in-series with muscle. Although these in-series elastic elements—e. g. tendons in vertebrates, or apodemes in invertebrates—can neither generate force nor do work, they are thought to bring substantial benefits to musculoskeletal performance; the mechanical properties of tendons, crucial determinants of these benefits, have consequently been subject of a large body of work. In sharp contrast, scarce information exists on the mechanical properties of apodemes. The little data that do exist appear to suggest that apodemes differ so substantially from tendons that their functional significance may differ, too. To increase our understanding of apodeme function, we determined the mechanical properties of the unguitractor apodeme (UTA) of *Sungaya aeta* stick insects. We devised an experimental protocol that permits tensile testing with slippery and brittle apodemes; we derived and validated a mechanical model that extracts the Young’s modulus from tensile tests with specimen with varying cross-sectional area, without the need for explicit measurement of the stress or strain distribution; and we interpreted the magnitude of the UTA modulus, strength and spring constant through allometric comparison with data on vertebrate tendons. The UTA modulus exceeds that of vertebrate tendons by almost one order of magnitude, but the size-corrected spring constant is nevertheless comparable and if anything smaller, due to systematic differences in apodeme and tendon shape. This many-to-one mapping suggests that apodemes may well convey the same functional benefits as tendons, and should not be prematurely excluded from invertebrate musculoskeletal models.

---

Movement is integral for bilateral animals, and driven by a highly conserved molecular motor: muscle (Steinmetz et al., 2012). A muscle’s mechanical output is limited by the force it can exert across different shortening lengths and velocities (Alexander, 2003; Biewener and Patek, 2018; Bobbert, 2013; Labonte, 2023; Labonte and Holt, 2024; McMahon, 1984; Sutton et al., 2019), and these physiological and mechanical limitations constrain animal performance, behaviour, and evolution (Labonte et al., 2024; Mendoza et al., 2023; Polet and Labonte, 2024). Muscle action is transmitted to the environment via skeletal elements, but rarely do muscle and skeleton connect directly. Instead, muscle action is mediated via tendons, aponeuroses (in vertebrates), or apodemes (in invertebrates). Although these elements are non-contractile—and thus can neither generate force nor do mechanical work—their presence is thought to bring great functional benefits to musculoskeletal performance (Alexander, 2002; Alexander et al., 1977; Biewener and Roberts, 2000; Holt and Mayfield, 2023; Roberts, 2016; Roberts and Azizi, 2011; Wilson and Lichtwark, 2011).

During running, energy that would otherwise be lost as heat can be transiently stored in form of elastic strain energy in stretched tendons, reducing the need for muscle work, and so presumably the expense of metabolic energy (Alexander, 1984, 1991; Alexander et al., 1982; Bennett and Taylor, 1995; Holt and Mayfield, 2023; Lichtwark and Wilson, 2005, but see Holt et al. (2014)); upon impulsive loading, tendons may act as shock absorbers that protect muscle fibres from critical damage (Dick et al., 2021; Konow et al., 2011; Roberts and Azizi, 2010; Roberts and Konow, 2013); tendons can significantly enhance muscle mechanical output during explosive movements, because the speed of elastic recoil is not restricted by the otherwise limiting force-velocity relationship of muscle (Aerts, 1997; Astley and Roberts, 2012; Galantis and Woledge, 2003; Labonte and Holt, 2024; Lichtwark and Wilson, 2005; Peplowski and Marsh, 1997; Richards and Sawicki, 2012; Sutton et al., 2019); and the rapid elastic response of tendons can passively stabilise movements against unpredictable perturbations, on time scales much faster than achievable via the nervous system, and so simplify necessary control strategies (Blickhan et al., 2007; Ghigliazza et al., 2005; Van Der Krogt et al., 2009). Without doubt, placing appropriately tuned elastic elements in-series with muscle can have diverse and significant benefits (Alexander, 2002; Alexander et al., 1977; Biewener and Roberts, 2000; Holt and Mayfield, 2023; Labonte and Holt, 2022; Roberts, 2016; Roberts and Azizi, 2011; Wilson and Lichtwark, 2011). The mechanical properties of *vertebrate* tendons have consequently been the subject of a substantial body of work. Perhaps surprisingly, much less is known about the mechanical properties of the invertebrate analogue: apodemes. The literature contains tendon data for upwards of 40 vertebrate species, covering a large range of body sizes, clades and ecological niches (Bennett et al., 1986; Bennett and Stafford, 1988; Ker, 1981; Lieber et al., 1991; Mossor et al., 2020; Pollock and Shadwick, 1994a,b; Shadwick, 1990; Shadwick et al., 2002, for a review, see Summers and Koob (2002)). In sharp contrast, only two reports on the mechanical properties of invertebrate apodemes exist to the best of the authors’ knowledge—both on the same species, and both published about half a century ago (Bennet-Clark, 1975; Ker, 1977).

In classic work, Henry Bennet-Clark sought to deter-mine the mechanical properties of the hindleg extensor tibiae apodeme of *Schistocerca gregari* locusts (Bennet-Clark, 1975). To this end, Bennet-Clark devised a cantilever-style bending experiment: small apodeme test pieces were pushed against a calibrated spring, resulting in apodeme deflection and spring compression. Apodeme samples were then fractured close to their fixed end, their thickness and width measured, and their elastic modulus estimated via standard beam theory, helped by the assumption that test pieces resemble beams with constant rectangular cross-section. These experiments were not without difficulty: even specimens as short as 3 mm “tapered appreciably” along their long axis (Bennet-Clark, 1975, p. 66), and thus deviated from the assumption of a constant cross-sectional area. To deal with this problem— and the “considerable variability” it introduced (Bennet-Clark, 1975, p. 66)—Bennet-Clark mounted six test pieces each, in one of two opposite orientations, hoping that this experimental design may correct for the mismatch between experimental reality and theoretical assumption. A Young’s modulus estimate of 18.9 ± 8.9 GPa, averaged across the 12 measurements, was the result. Bennet-Clark also attempted to measure the apodeme’s tensile strength. Initial trials with isolated apodeme test pieces gave “low values” (Bennet-Clark, 1975, p. 56), and were not discussed further. Instead, Bennet-Clark attached a polyester sling around the tibia, cut the suspensory ligaments of the femoro-tibial articulation, clamped the femur, and then applied a load to the sling to pull the assembly apart. This method made it difficult to apply loads “symmetrically, smoothly and rapidly enough”, and, in many cases, the preparation broke with the apodeme still intact (Bennet-Clark, 1975, p. 66). These considerable difficulties cautioned Bennet-Clark—a famously gifted and thorough experimentalist—to report only an approximate range for the maximum load the apodeme can sustain—”probably not less than 14 N and not more than 17 N” (Bennet-Clark, 1975, p. 66)—roughly equivalent to an ultimate tensile strength between 0.56-0.68 GPa. Two years later, Robert Francis Ker reported mechanical property measurements on two apodemes of *S. gregari* locusts in his PhD thesis, completed under the supervision of AC Neville (Ker, 1977). Ker constructed a miniature tensile tester to measure the Young’s modulus of the tibial section of the unguitractor apodeme (in his thesis referred to as pre-tarsal flexor), as well as of the distal region of the tibial extensor apodeme (Ker, 1977). In the thesis abstract, Ker notes that “most of the [mechanical property] results are approximate” (Ker, 1977, p. i), a critical self-assessment best understood by revisiting the details of his experiments. Ker tested apodemes in tension via the application of dead loads. For the unguitractor apodeme, he extracted a Young’s modulus estimate of 15 GPa from *one* apodeme tested in liquid paraffin (to limit water loss) at three different lengths (to correct for “end effects”. See section 2.4.3 in Ker, 1977, for further detail). Modulus estimates of 13, 13 and 11 GPa resulted from three further tests with two apodemes. In two of these tests, the modulus was estimated from the extension at *one single* dead load; two apodemes were submerged in water, and one in liquid paraffin. The overall average modulus was thus 13 ± 1.73 GPa, obtained from three independent samples. Ker himself notes that “the number of measurements is insufficient to make any firm statements regarding the variation of Young’s modulus” (Ker, 1977, p. 54), and attributes much of this limitation to the difficulty in area and length measurements for small specimen. He then proceeded to use the same protocol to measure the modulus of the distal region of tibial extensor apodeme, selected because “irregular variations in area are measurable but are not sufficient to have any serious consequence” (Ker, 1977, p. 58). Ker reported a modulus of 8.1 GPa for *one* single apodeme (tested in liquid paraffin); three further tests were conducted with specimen ultimately deemed “too short to give a reliable value” (Ker, 1977, p. 64); and two final tests resulted in modulus estimates of 9.75 GPa (in liquid paraffin), and 11.4 GPa (in water). Thus, the average Young’s modulus came to 9.75 ± 1.65, estimated from three independent samples. Ker summarised his results with great caution: “the Young’s modulus of *a particular apodeme* has been found to be 8.1 GPa. Incomplete evidence indicates that this is a low value within the range that may occur.”(emphasis added, Ker, 1977, p. 81). As Bennet-Clark before him, Ker also attempted to measure the apodeme’s ultimate tensile strength, but he, too, had to give up: results were “very variable” (Ker, 1977, p. 62), and the sustained loads were never high enough “to be reasonable” (Ker, 1977, p. 61).

Revisiting the pioneering work of Bennet-Clark and Ker in detail provides three valuable lessons.

First, measurements of apodeme mechanical properties there are few, and those that do exist are subject to considerable uncertainty: the sample size was small, the experimental difficulties large, and two independent modulus estimates for the same apodeme in the same species differ by more than a factor two (8.1 GPa in Ker (1977) vs 18.9 GPa in Bennet-Clark (1975). We note that the two studies did not test the exact same apodeme section, so that this difference may reflect a true variation in modulus along the apodeme long axis). This uncertainty is a just cause for concern, for a significant body of subsequent work relied on these reports, be it to inform musculoskeletal modelling (Cofer et al., 2010; Full and Ahn, 1995; Schmitt and Holmes, 2003), to attribute jump performance to specific leg “springs” (Gabriel, 1985; Rogers et al., 2025, 2016), to compare apodeme and tendon safety factors (Alexander, 1981), to determine leg muscle strain and function (Ahn and Full, 2002; Ahn et al., 2006; Full et al., 1998), to assess the “tuning” between muscle and apodeme properties (Rosario et al., 2016; Sutton et al., 2019), or, qualitatively, to discuss differences in the invertebrate vs vertebrate flight apparatus (Wold et al., 2023).

Second, the uncertainty in the original work—as much as the absence of later work that reports modulus or strength data—is not without good reason: testing apodemes in tension is challenging, as “only the insect is able to make attachments to it which do not cause it to break” (Brown, 1963, as cited by Bennet-Clark (1975)); modulus measurements by any means—bending, tension or compression—are complicated further by the variation of the apodeme’s cross-sectional area along its long axis (e. g. Bennet-Clark, 1975; Gabriel, 1985; Ker, 1977). One way out of this conundrum is to extract apodeme test pieces so short that the variation in cross-sectional area remains small—but clamping becomes more challenging still, and length measurements so inaccurate that labour-intense “free length” corrections (or technically involved direct strain measurements) become necessary (Ker, 1977). Another route is to attempt to average out cross-sectional area variation by mounting specimen in variable orientations for bending tests (Bennet-Clark, 1975); but the error introduced by this experimental approach is unclear. To the best of our knowledge, a suitable methodological approach that helps resolve these problems remains absent, and with it more robust estimates of the mechanical properties of apodemes. Consequently, our understanding of apodeme function in invertebrates is stuck far behind what we know about the function of tendons in vertebrates.

Third, and on the note of comparison, the limited information that does exist suggests that the modulus of apodemes may be more than one order of magnitude larger than that of tendons. This difference matters, because it suggests that the benefits of in-series elasticity may differ between clades, too. In-series elastic elements derive much of their acclaimed functional benefits from their ability to stretch and recoil by neither too much nor by too little under physiological loads (Alexander, 2002; Alexander et al., 1977; Biewener and Roberts, 2000; Holt and Mayfield, 2023; Roberts, 2016; Roberts and Azizi, 2011; Wilson and Lichtwark, 2011): very stiff tendons may stretch only little, and eventually become functionally irrelevant altogether; overly compliant tendons, in turn, will prevent muscle from developing significant force, and only store negligible strain energy. So how stiff are invertebrate apodemes in comparison to vertebrate tendons, and what does a potential difference imply about their function?

To begin to answer this question, and to provide a suggestion for how the highlighted methodological and theoretical difficulties may be resolved, we here report measurements of the elastic modulus, tensile strength, and spring constant (stiffness) of the unguitractor apodeme of *Sungaya aeta* Hennemann, 2023 (Phasmatodea: Heteropterygidae) stick insects. These measurements are enabled by a short theoretical development that permits measurement of the modulus and prediction of the spring constant of test pieces with arbitrary cross-sectional area variation, and supplemented by a brief comparative analysis of the allometry of the mechanical properties of in-series elastic elements.

## Materials and methods

### Study animals

Adult female *Sungaya aeta* Hennemann, 2023 (Phasmatodea: Heteropterygidae) stick insects were used for all experiments (n = 13, body mass 3946 ± 678 mg); we chose this species for its general availability, relatively large body size, and ease of husbandry. Individuals were taken from a laboratory colony maintained in a 12:12 h day and night cycle at 25 ^°^C, 60% relative humidity, housed within a climate chamber (FitoClima 12.000 PH, Aralab, Rio de Mouro, Portugal). Bramble, replaced weekly, was provided as *ad libitum* food source. Individuals were sacrificed through freezing for at least 30 min, and subsequently weighed to the nearest 1 mg (OHAUS Explorer analytical balance EX124, max. 120 g *×* 0.1 mg, Parsippany, NJ, USA).

### Apodeme extraction and storage

Specimens were thawed for 10 min prior to apodeme extraction. We decided to characterise the mechanical and morphological properties of the unguitractor apodeme (UTA), first, because it runs continuously through the femur, tibia and tarsus to actuate the tarsomeres and claws, and is thus of considerable length, and second because it is a general feature of the insect limb anatomy (Figure 1 a. See Ichikawa et al., 2016; Radnikow and Bässler, 1991). To extract UTAs, the right hind legs, chosen arbitrarily, were amputated at the trochanter-femur joint using a scalpel. To ensure that the UTA and its attachment to the unguitractor plate remained intact, we completed the next step under a Leica SAPO stereo microscope (Zill et al., 2014): a pair of micro scissors was used to pierce through the exoskeleton, and to create an incision along the pretarsal claws; a gentle pull on the claws with tweezers was then sufficient to dislodge the UTA from its proximal attachment site, making it possible to pull it out of the leg in its entirety with little force. Extracted UTAs were either immediately subjected to imaging and tensile testing (see below), or stored at approximately -20 ^°^ C for no more than three days, a preservation method most likely to maintain cuticle mechanical properties (Aberle et al., 2016). Prior to and throughout all experiments, UTAs were kept at ambient conditions, and sprayed with pH 7.4 phosphate-buffered saline (PBS) every 30 s to keep them hydrated (Bennett et al., 1986; Ker, 1981; Klocke and Schmitz, 2011; Mossor et al., 2020). Visual inspection confirmed that no salt crystals formed on the apodemes throughout the measurement time.

**Figure 1.**
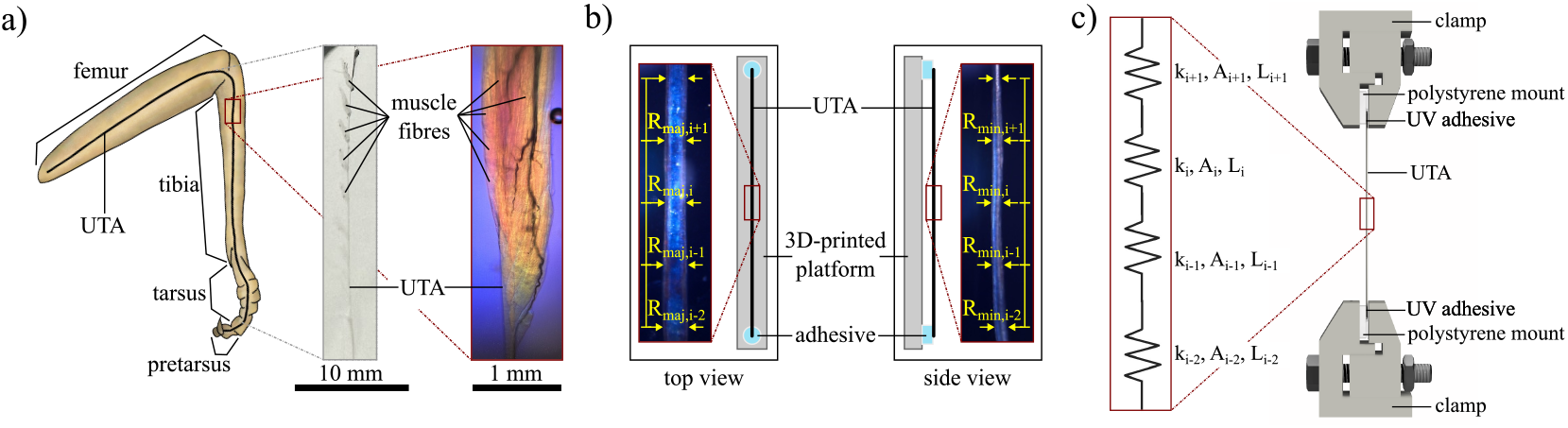
a) A common feature of animal legs are elastic elements that connect limb muscle with the skeletal elements they actuate. In insects, one prominent such elastic element is the unguitractor apodeme (UTA), which runs from its origin in the proximal section of the femur all the way through the tibia and tarsus, where it connects onto the unguitractor plate in the pre-tarsus. Muscles attach to this apodeme in both the femur and tibia, and control tarsal segments, as well as the pre-tarsal attachment system (Endlein and Federle, 2008; Federle et al., 2004; Ichikawa et al., 2016; Radnikow and Bässler, 1991). b) The UTA cross-sectional area varies along the UTA length, rendering mechanical characterisation challenging. To quantify this variation, we approximated the UTA cross-sectional area as an ellipse characterised by a major and a minor axis, *R*_maj_ and *R*_min_, respectively, informed by preliminary inspection via light microscopy. The major axis is approximately orthogonal to the dorsal-ventral axis of the leg, and its magnitude differs most significantly from the minor axis where the UTA crossed from the femur into the tibia. To quantify both axes across the UTA length, UTAs were glued onto a sample stage that enabled consistent rotation by 90^°^, permitting the acquisition of two orthogonal light microscopy images, and subsequent measurements of both radii via Fiji v2.14.0. c) Accounting for the variation in UTA cross-sectional area along the sample length is necessary for accurate detailed mechanical characterisation: because the cross-sectional area varies along the sample length, so do stress and strain. To appropriately model this variation, the UTA was discretised into *i* segments with local cross-sectional area *A*_*i*_ and spring constant *k*_*i*_, each approximately 500 µm in length (for details, see text). To test UTAs in tension, they were affixed to roughened polystyrene sheets using UV glue, clamped, and strained until failure with an average strain rate of 0.0014 ± 0.000098 s^-1^.

### Apodeme morphometry

Preliminary inspection of test samples via light microscopy revealed that the UTA cross-section was not uniform. Instead, it can be approximately described by an ellipse, defined by a minor and a major axis, *R*_*min*_ and *R*_*maj*_, which both vary along the apodeme long axis (see also images of apodeme cross-sections in Ker, 1977, p. 49 and 59). *R*_*maj*_ is approximately orthogonal to the dorsal-ventral axis of the leg, and its magnitude differed most significantly from *R*_*min*_ where the UTA crossed from the femur into the tibia (see results); in this region, the UTA is significantly wider than it is thick. The UTA major and minor axes were consequently defined as the two axes aligned with and perpendicular to this widest crosssectional diameter, respectively. To capture the variation of *R*_*min*_ and *R*_*maj*_ quantitatively, a simple light microscopy protocol was developed (Z6 macroscope, Leica, Wetzlar, Germany). The proximal and distal ends of each UTA were first adhered to a 3D-printed pentagonal prism (55 mm length *×* 10 mm height *×* 5 mm base width), using a thin strip of blue tack (Figure 1b). The pentagonal shape of the prism helped minimise obstruction of the UTA during lateral imaging, and provided orthogonal faces, which facilitated consistent rotation between perpendicular views. Prior to attachment, the UTA was straightened by suitable rotation as necessary, such that one image plane was approximately perpendicular to the dorso-ventral axis and *R*_*maj*_. High-resolution images were then captured along the UTA long axis (Figure 1 b), utilising the native motor stage control routines implemented in the microscope software (Leica LAS X). The pentagonal prism was subsequently rotated by 90^°^ onto an adjacent face, and the imaging process was repeated to capture a second, orthogonal view of *R*_*min*_. UTA images were then imported into Fiji v2.14.0 (Schindelin et al., 2012), to manually measure *R*_*min*_ and *R*_*maj*_ at approximately 500 µ m intervals, yielding 85 cross-sectional area measurements per UTA on average. The local cross-sectional area at these points, *CSA*_local_, was estimated from these measurements as that of an ellipse with axes *R*_*min*_ and *R*_*maj*_, *CSA*_local_ = *πR*_*min*_*R*_*maj*_—the UTA was thus effectively discretised along its long axis into 500 µm sections, each characterised by a constant local cross-sectional area *CSA*_local_ (Figure 1 b).

To extract a single representative variation of *R*_*min*_, *R*_*maj*_ and *CSA*_local_ across the UTA long axis from all 13 independent measurements, we conducted an arc-length analysis without signal registration in ARCGen 2024.1.0 (Python v3.9.13), a technique developed specifically to provide appropriate averages for bivariate biomechanical data that vary in length (Hartlen and Cronin, 2022).

### Mechanical characterisation

#### Determining the elastic modulus and spring constant of samples with arbitrary variations in cross-sectional area

Three biomechanical properties of functional relevance for tendons and apodemes are their ultimate tensile strength—the stress at rupture, *σ*_max_ = *F*_*max*_*/A*; their Young’s modulus, *E*—the proportionality constant between stress and strain for small strains *E* = *σ/ε*; and their spring constant or stiffness, *k*—the proportionality constant between force and displacement for small displacements, *k* = *F/δ* = *EAL*^−1^, where *A* and *L* are the relevant cross-sectional area and length, respectively. All three properties are readily determined from force-displacement data if the sample cross-sectional area is known and constant throughout the sample length. If the cross-sectional area is variable, however, a problem arises, for the distribution of stress and strain in the sample is now non-uniform, too. What is the appropriate definition of strength, modulus and spring constant in samples with non-constant cross-section and thus spatially variant stress and strain?

A sample will fail in tension if the maximum stress it experiences exceeds its ultimate tensile strength. The maximum stress occurs in the cross-section with the smallest area, and we thus extracted *σ*_max_ as the ratio between maximum force and the minimal cross-sectional area in the unclamped “free” sample region:

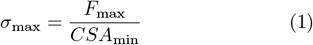

Sample inspection *in post* confirmed that failure typically occurred in apodeme regions with small crosssectional area.

To estimate the Young’s modulus, we will now derive an equivalent (representative) single-valued stress and strain, such that *E* = *σ*_eq_*/ε*_eq_. In general, these equivalent parameters are not unique, and the only condition on them is that their ratio equals the “true” modulus *E*. Our specific choice is pragmatic in the sense that it permits to infer *σ*_*eq*_ and *ε*_*eq*_—and thus *E*—without the need to explicitly characterise the spatial variation of strain or stress. Instead, we rely on the operational definition of strain in classic tensile tests, which remains straightforward to use, no matter the variation of sample cross-sectional area:

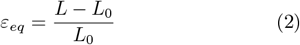

Here, *L* and *L*_0_ are the length of the free (unclamped) sample region for the deformed and undeformed sample, respectively.

Consider, now, a sample comprising *i* discrete segments in series, each with identical elastic modulus *E*, but variable undeformed length *L*_*i*,0_ and cross-sectional area *A*_*i*_ (Figure 1 c). The undeformed sample length, *L*_0_, is simply the sum of the individual segment lengths, *L*_0_ = *L*_*i*,0_. Upon application of a force *F*, each section stretches to a new length, *L*_*i*_, which follows from Hooke’s law as:

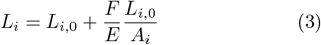

where we relied on the fact that elements in-series experience the same load. The equivalent (globally measured) strain thus reads:

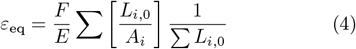

Upon insertion of *A*_*i*_ = *F/Eε*_*i*_, this particular definition of the equivalent strain, *ε*_eq_, may be identified as the arithmetic mean strain, 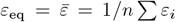. Combining this result with the definition of the elastic modulus yields the appropriate definition for the equivalent crosssectional area that determines the equivalent stress:

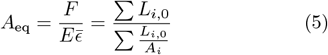

which—remarkably—corresponds to the weighted harmonic mean area, *A*_eq_ = *Ã*. The elastic modulus of an otherwise homogenous sample with arbitrary variations in cross-sectional area may thus be defined as the proportionality constant between the weighted harmonic mean stress and arithmetic mean strain:

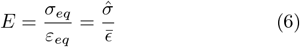

For a sample with constant cross-sectional area, eq. 6 reduces to the standard definition, for the arithmetic and the weighted harmonic mean area are now equal. By replacing the finite sum over *i* discrete elements with a sum over an infinite number of infinitesimal terms (an integral), eqs. 4 and 5 can of course also be written as exact continuous expressions—but this requires knowledge of an integrable function, *A*_*i*_(*L*), generally unavailable for biological samples with complex and irregular geometry; some form of discretisation is thus often unavoidable. We close this short derivation by highlighting the assumption that the modulus is constant throughout the sample length, arguably parsimonious, but nevertheless unverified.

The apodeme modulus is a material property and thus of considerable interest. It is, however, not the functionally relevant mechanical quantity, for it is only one of three parameters that define the apodeme’s spring constant, i. e. the proportionality constant between force and displacement. In contrast to *E, k* is a system property that depends on material properties as much as on the apodeme geometry, *k* = *EAL*^−1^. As for the modulus, the question arises what the correct definition for *A* may be when sample cross-section is irregular. The above derivation at once identifies the weighted harmonic mean area as appropriate choice:

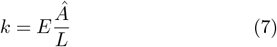

However, the difficulties are not yet over, because *Ã* depends on the apodeme section that provides the length *L*. Extracting *k* directly from a tensile test may seem like the simplest choice, but will yield erroneous estimates, for the gauge length will typically be different from the effective free length of the apodeme *in vivo*. This effective free length, *L*_*f*_, is not trivial to determine; in fact, there are multiple rational definitions of *L*_*f*_, because muscles attach to the UTA at several points along its length (Radnikow and Bässler, 1991). To maintain generality, we assess the variation of UTA spring constant for any possible choice for *L*_*f*_, *L*_*f*_ = *νL*_0_, where 0 *< L*_0_ ≤ 1: we evaluate *Ã*(*ν*), and then estimate the spring constant *k*(*ν*) via eq.7. A representative spring constant is once again extracted from individual data via an arc length analysis, this time conducted on log10-transformed spring constant to account for the fact that *k* diverges as the length *L* approaches zero. A more detailed assessment of apodeme spring constant, informed by a comparative analysis and musculoskeletal anatomy, is provided in the discussion.

To the best of our knowledge, the definition of a modulus and spring constant for a sample with non-uniform cross-section has not been discussed in previous work on tendons or apodemes, which used the operational (global) definition of sample strain without exception. Perhaps the variation in sample cross-sectional area was deemed small enough to be negligible—this aspect will be addressed in more detail in the discussion.

#### Model verification

To verify eq. 6, we performed validation experiments on a mechanically well-defined elastomer, polydimethylsiloxane (PDMS; SYLGARD 184, Dow, Midland, MI, USA). Two sets of PDMS samples were cast (n = 5 for each): they shared the same mixing ratio and curing protocol (10:1 silicon base:curing agent; 4 h at 65 ^°^C), but differed in geometry. The first sample set had a length of 100 mm (excluding a broadened region for gripping, Figure 2), and a constant cross-sectional area (area = width *×* thickness; 24.12 mm^2^ = 8.04 *×* 3.00 mm). The second set of samples had the same free length and an almost identical arithmetic average CSA (24.01 mm^2^), but the local CSA varied considerably along the sample length, in a manner which roughly resembled the representative variation determined for the UTAs via the arc length analysis (see above). To obtain a parametrised form of this variation, the representative CSA curve was fitted to a fifth order polynomial, the lowest order that met two criteria: (I) the ratio between maximum and minimum crosssectional area, as predicted by the polynomial, was similar to the ratio extracted from the arc-length analysis (17.65 vs. 17.80, respectively, see Figure 2 a)); and (II), the expected ratio of spring constants between a sample with variable cross-sectional area and a sample with the same arithmetic mean but constant cross-sectional area, mirrored the expectations from the arc-length analysis (0.529 vs 0.531, respectively). Based on the fitted function, we designed sample geometries in Inventor v2024 (Autodesk Inc. San Francisco, CA, USA), and then 3D-printed PETG moulds (MK4, Prusa Research, Prague, Czech Republic), from which sample sets with either variable or constant cross-sectional area were fabricated (Figure 2 b): moulds were filled with uncured but degassed PDMS, covered with a silanised glass plate clamped on top, and put into an oven for curing as described above.

**Figure 2.**
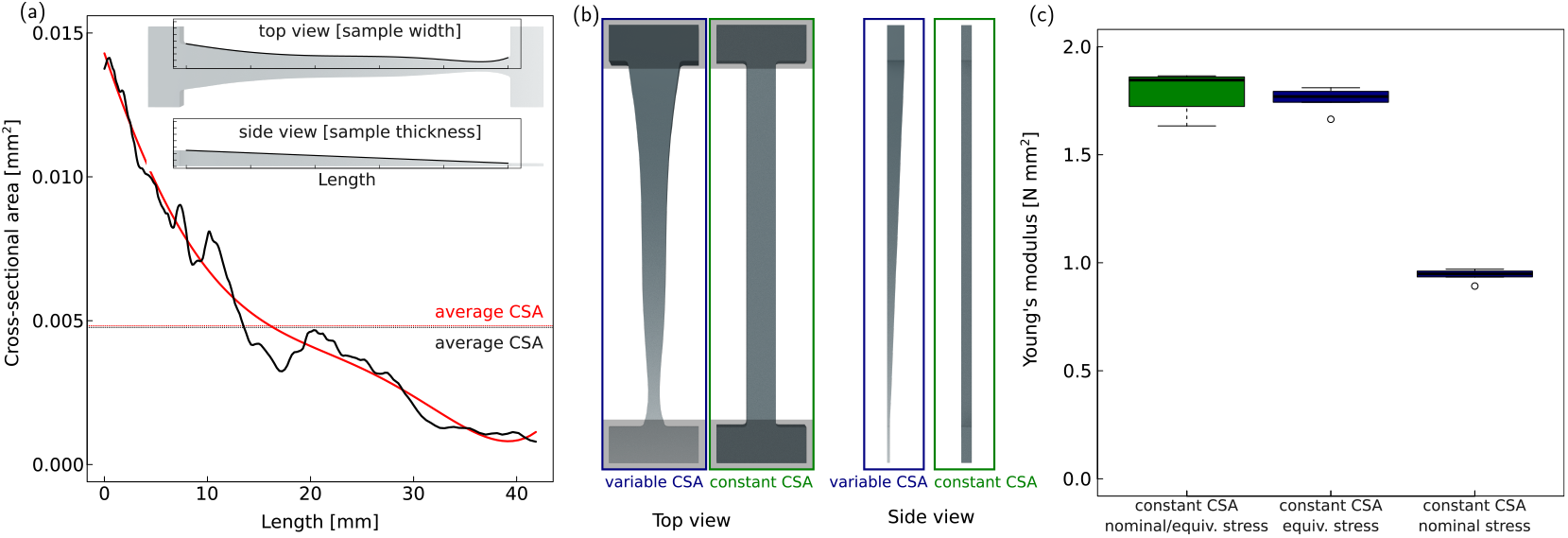
Extracting the Young’s modulus of apodemes via tensile testing is challenging, due to the considerable variation in cross-sectional area along the apodeme long axis. To solve this problem, we derived an equivalent stress and strain, such that their ratio equals the Young’s modulus regardless of specimen shape (see text). To verify this derivation, two sets of PDMS samples were fabricated. (a) The first set resembled the cross-sectional area variation observed for apodemes, as characterised by a 5^th^ order polynomial fit (red) to an arc-length analysis on the apodeme raw data (black; see text and Figure 4); (b) the second set had the same arithmetic mean cross-sectional area, but this time this area was constant throughout. Both samples were fabricated using 3D-printed moulds and equal curing conditions, before submitting them to tensile testing at a strain rate of 0.00167 s^-1^ (n=5 per set). The global or arithmetic average strain, 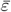 was used as the equivalent strain because it can be easily extracted from the global displacement data; the modulus was then extracted using either the arithmetic average or weighted harmonic mean stress, 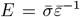 or 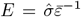, respectively. (c) Although both sample sets were made from the same material, the moduli extracted via the arithmetic mean stress differed by about a factor two between the two samples. In contrast, using the weighted harmonic mean stress for both samples yielded modulus estimates that were statistically indistinguishable, verifying the derivation of the equivalent stress and strain. Note that other definitions for the equivalent stress and strain, while possible, are perhaps less desirable, because they will require direct measurements of the strain or stress distribution across the sample.

Both sample sets were then tested in tension using a MultiTest 5-xt (Mecmesin, Slinfold, UK; 25 N load cell), a gauge length of 100 mm, and a constant speed of 10 mm min^−1^, corresponding to a strain rate of 0.00167 s^−1^. Prior to testing, the clamping distance was measured to the nearest 0.01 mm with a digital calliper (SEN-331-1230K, The Senator Group, Accrington, UK). Stress-strain curves were constructed from the force-displacement data using the global or arithmetic mean strain 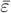, and either the arithmetic or weighted harmonic mean cross-sectional area, 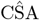 or CŜA, respectively; the Young’s modulus was then extracted as the slope of an ordinary least-squares regression between 0-3% of the equivalent strain. We note that PDMS remains linearly elastic across a larger strain range (0-10%, Johnston et al., 2014; Palchesko et al., 2012). However, due to the local variations in CSA in one sample set, the local strain exceeds the equivalent strain, so that equivalent strains below 3% were necessary to ensure that the local strain did not exceed 10% anywhere in the sample.

The Young’s modulus extracted for the control sample set with constant CSA was 1.79 ± 0.10 MPa, in robust agreement with independent measurements for PDMS samples with equal mixing ratio (Johnston et al. (2014), Figure 2 c). For the sample set that mimicked the CSA variation of UTAs, the modulus based on the arithmetic mean stress, 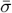, was 0.94 ± 0.03 MPa, almost half of what was measured for the control samples. Using the weighted harmonic mean stress, 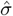, however, yielded a modulus of 1.76 ± 0.06 MPa, statistically indistinguishable from the control measurement (two sample t-test: *t*_8_ = 0.56, p = 0.59).

#### Apodeme tensile testing

Having validated a method to obtain the Young’s modulus from samples with non-uniform cross-section, we were in the position to evaluate the Young’s modulus and ultimate tensile strength of the UTA via uniaxial tensile tests. A total of 13 samples were tested using an ElectroForce ®3200 Series III (New Castle, Delaware, USA), equipped with screw gauge clamps (Figure 1c).

Much like earlier attempts to test apodemes in tension, we encountered experimental difficulties, and unsuccessfully tested a series of mounting options, summarised briefly here for the benefit of future experimenters. In an attempt to ensure that minimum clamping pressure suffices to secure wet and thin apodemes, we initially sought to fixate them between form-fitting materials, such as polydimethylsiloxane sheets or multiple types of sticky tape (Iatridis et al., 2003). However, no material sandwich was able to prevent UTA slippage, most likely due to the formation of air bubbles in the assembly during mounting. To increase clamp grip, we next followed recommended practice and attached sandpapers of different grit sizes to the clamps (Jiang et al., 2020). Unfortunately, sample slippage still occurred for the majority of samples. Previous work on mammalian tendons avoided these issues through the use of “winding clamps” (Grant et al., 2015): tendons were wrapped around an elongated cylindrical rod to fixate them via the Capstan effect. However, it proved impossible to wrap stiff apodemes without fracturing them. By trial and error, we eventually identified a combination of the first two approaches that worked for the vast majority of preliminary test samples, and was consequently used as preparation method for the final set of experiments: 10 *×* 5 *×* 2 mm polystyrene sheets were roughened with sandpaper to increase grip, and approximately 4 mm of each UTA were affixed to them on both ends using UV glue (Venaird Bug Bond Lite Fly Tying UV Resin, London, England). The glue was cured by UV irradiation for 15-20 s, and the samples were fixed into the gauge clamps (Figure 1 c), such that the average free sample length was 35.26 ± 2.51 mm, as determined again using a digital calliper.

A preload of 0.005 N—equivalent to approximately 0.1 % of the tensile force at rupture as determined in preliminary experiments—was applied to remove any UTA slack, and the samples were subsequently extended at a constant speed of 0.05 mm s^−1^ until failure. The low testing speed was chosen to limit the average strain rate to 0.0014 ± 0.000098 s^−1^, ensuring approximately quasi-static loading conditions, and in doing so avoiding further complexity due to sample viscoelasticity (which is likely minimal to begin with, Ker, 1977). Tested samples were reimaged under the Z6 macroscope to verify that apodeme failure had occurred away from the glue interface, and was thus not driven by stress concentrations close to the clamping site.

Equivalent stress-strain curves were then extracted using the weighted harmonic mean stress and arithmetic mean strain. The arithmetic mean strain was calculated from the free sample length and gauge displacement as per eq. 2, and the weighted harmonic mean area was estimated from the light microscopy data for the free sample region via eq. 5; dividing the measured load by this area yielded the equivalent stress. The elastic modulus was then extracted as the gradient of the relationship between weighted harmonic mean stress and arithmetic mean strain, estimated via ordinary least squares regression in the region 0.002 *< ε*_eq_ *<* 0.007 (Figure 3).

**Figure 3.**
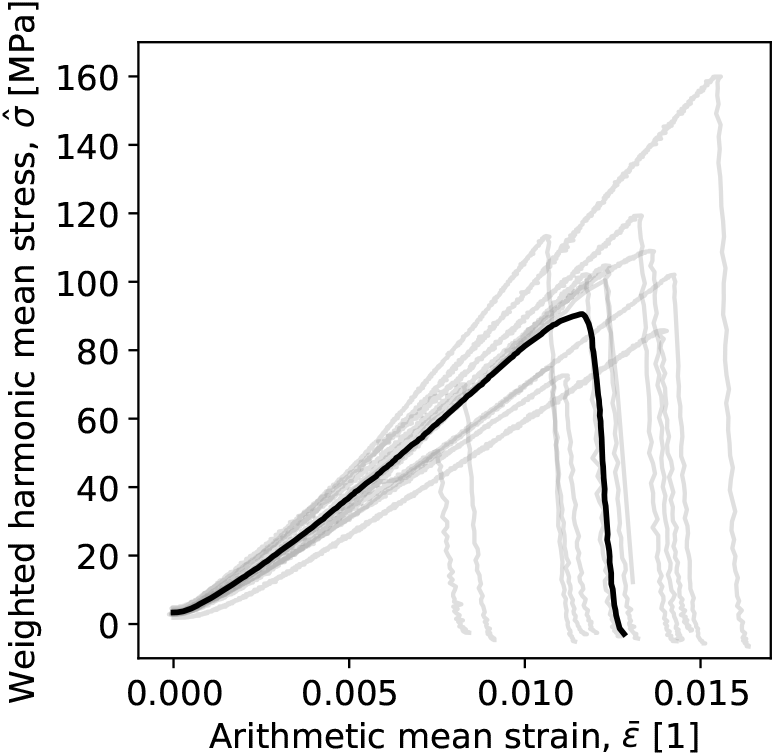
The Young’s modulus of unguitractor apodemes was extracted as the ratio between the weighted harmonic mean stress and arithmetic mean strain, 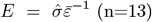. The J-shape response characteristic of Mammalian tendons was absent, and apodemes failed catastrophically once a critical force was reached, indicating brittle material behaviour. The solid line is a representative stress-strain curve, extracted from the raw data via an arc-length analysis (Hartlen and Cronin, 2022). Note that the plot does not start from [0, 0] as a pre-load was applied to remove any specimen slack, and that maximum stress and strain in the sample are larger than the maxima that can be read from this plot, due to the variation in cross-sectional area across the length of the test pieces.

## Results

### Apodeme morphology

UTAs had an average length of 42.89 ± 3.67mm (n=13). Both *R*_*min*_ and *R*_*maj*_ varied considerably along the UTA long axis; global maxima of about 0.15 and 0.12 mm occurred close to the attachment site onto the unguitractor plate; global minima of about a quarter to a third of these values occurred at the UTA’s proximal end, respectively (Figure 4 a-b). *R*_*min*_ decreased from distal to proximal, reaching close to its minimum magnitude by the mid-point (Figure 4 a-b). The decrease in *R*_*maj*_ initially mirrored the decrease in *R*_*min*_, but it then markedly widened in the section that spans the femur-tibia joint in the proximal half of the UTA. In this section, extending over about a quarter of the total UTA length, *R*_*maj*_ first increased to a local maximum of 0.1 mm, about double the local magnitude of *R*_*min*_, before it declined again to reach its global minimum of about 0.04 mm (Figure 4 a-b). Resulting from the marked variation in *R*_*min*_ and *R*_*maj*_ was a substantial variation of the local cross-sectional area, CSA_local_ (Figure 4 c). Close to the unguitractor plate, CSA_local_ exhibited a global maximum of CSA_*local*_ ≈ 0.015 mm^2^. It then decreased steeply to about a quarter of this value, rose to a local maximum of 0.005 mm^2^ in the section that spans the femur-tibia joint, and eventually dropped to a global minimum of about 0.002 mm^2^ at its proximal end. This variation in cross-sectional area resulted in a marked difference between the arithmetic and weighted harmonic mean cross-sectional areas, 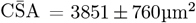 vs CŜA = 1735 ± 516µm^2^, respectively (n=13; Figure 5 a).

**Figure 4.**
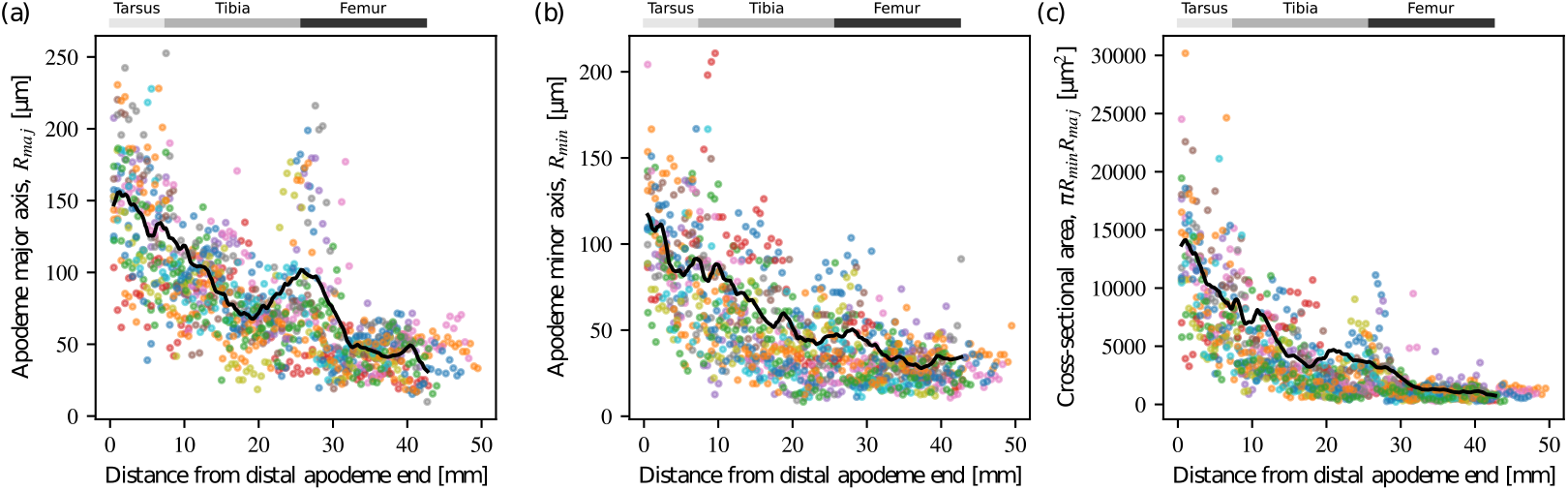
Apodeme cross-sectional area varied considerably along the apodeme’s long axis. To characterise this variation, the apodeme cross-section was approximated by an ellipse with a major and minor axis, *R*_maj_ and *R*_maj_, defined as the two axes aligned with and perpendicular to the widest apodeme axis at the point where it crosses from the femur into the tibia, respectively. Both radii varied substantially across the apodeme length (a and b)—the apodeme was widest and thickest close to the insertion site onto the unguitractor plate, and thinnest in the proximal femoral section. Local maxima appear where the apodeme crosses joints, and are most pronounced at the transition between the tibia and the femur. As a result of the radius-variation, the cross-sectional area varied substantially too: the smallest and largest cross-sectional area differed by almost one order of magnitude. The solid black line indicates a representative variation, extracted from the raw data via an arc-length analysis (Hartlen and Cronin, 2022); grey bars on the top indicate the average length of major limb segments for a hypothetical UTA with a length of 42 mm, corresponding to the UTA modelled by the arc-length analysis. n=13 for all plots, with sample ID indicated by colours.

**Figure 5.**
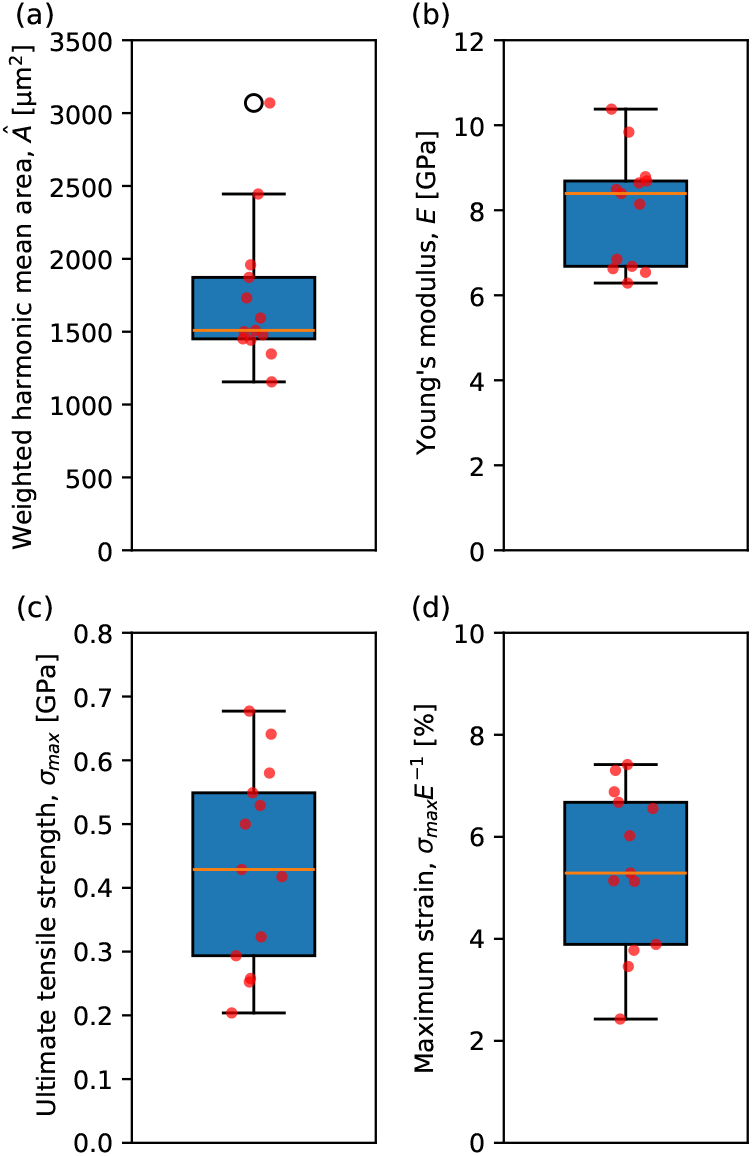
Boxplots summarising results for (a) the weighted harmonic mean cross-sectional area; (b) the Young’s modulus, extracted as the ratio between weighted harmonic mean stress and arithmetic mean strain, 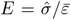; (c) the ultimate tensile strength, extracted as the ratio between maximum force and minimal cross-sectional area, *σ*_max_ = *F*_max_*/*CSA_min_; and (d) the maximum strain at failure, *ε*_max_ = *σ*_max_*/E*. Boxplots show the median, the first and third quartiles, and 1.5 *×*the interquartile range (n=13 for all data).

### Apodeme mechanical properties

UTA stress-strain curves were approximately linear, until a rapid drop in stress indicated abrupt and brittle sample failure (Figure 3, see also Ker, 1977, p. 51 for similar accounts for locust apodemes). The J-shaped stress-strain response characteristic of mammalian tendons was absent (Pollock and Shadwick, 1994b), and the data was substantially linear (*R*^2^=0.99 ± 0.001, n=13; Figure 3). This absence may be a result of local strains much in excess of the global strain, so that the linear response of strongly strained sections dominates the response for small global strains already; it should thus not be taken as strong evidence against small strain non-linear elasticity of the UTAs in general.

The elastic modulus of the UTA, estimated as the ratio of the weighted harmonic mean area and arithmetic mean strain, was *E* =8.03 ± 1.27 GPa (n=13). Modelling the UTA as a sample with a constant cross-section, equiv-alent to 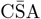, instead would yield a modulus estimate of *E* = 3.51 ± 0.44 GPa (n=13), or about half the best estimate of the true value, directly mirroring the difference between CŜA and 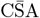 (see above). UTAs had a maximum tensile strength of *σ*_max_ = 434.85 157.95 MPa (n=13), about a twentieth of the modulus, *σ*_max_ ≈ *E/*20. The arithmetic mean strain at failure was 0.012 ± 0.002 (n=13), but the local strain in the failure region was almost four times higher, *σ*_max_*/E* ≈ 0.05, and thus close to critical strains reported for mammalian limb tendons (Pollock and Shadwick, 1994b).

Apodeme modulus and weighted harmonic mean crosssectional area are two of the three determinants of the apodeme’s spring constant—a system property that also explicitly depends on the apodeme’s free length, *L*_*f*_. The variation of spring constant with *L*_*f*_ is not trivial, because the weighted harmonic mean area is itself a function of *L*_*f*_, *Ã*(*L*_*f*_) (Figure 6). It is instructive to consider as reference the spring constant of a hypothetical apodeme with equal modulus, *E* = 8.1 GPa and total length, *L*_0_ = 42 mm, but a constant cross-sectional area, equal to the apodeme’s arithmetic mean area, *Ā* = 3851 µm ^2^. The free length is then the only variable, and the spring constant reads *k* = 30.1 NL^−1^. Such an apodeme thus has a minimum spring constant of 733 N m^−1^ when it deforms along its entire length; double this spring constant at half its length; and 10 times this spring constant at a tenth of its length. The spring constant of the real apodeme, which varies in crosssection, is about 300 N m^−1^, 1400 N m^−1^, and 20,000 N m^−1^ at the same lengths instead—it is half, about equal, and about three times as stiff; its spring constant thus varies much more rapidly (Figure 6).

**Figure 6.**
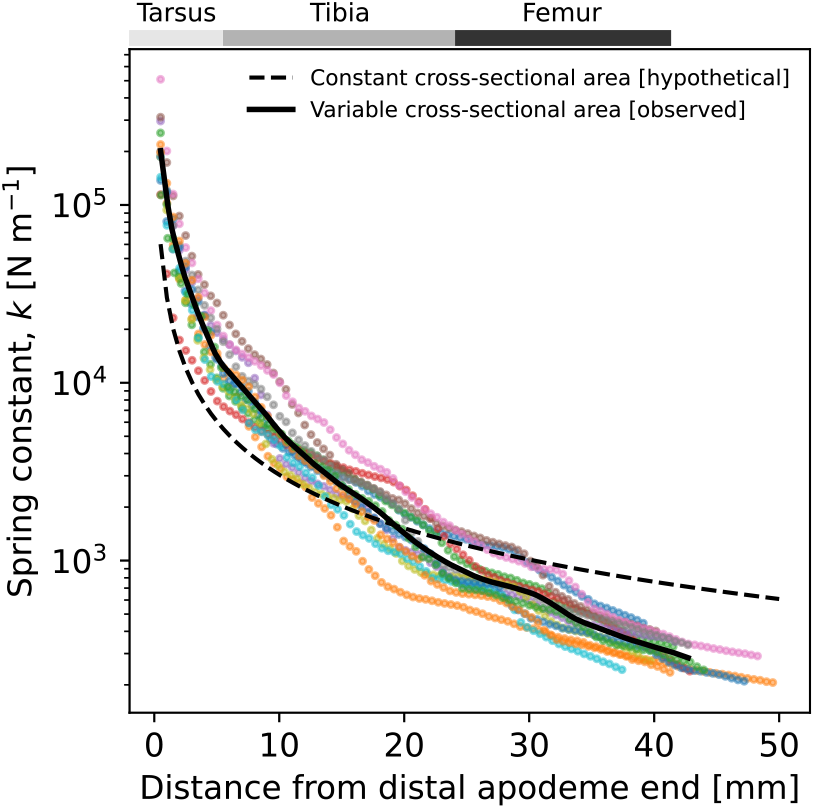
The spring constant varies with the sample free length, *k* ∝ *L*^−1^. Because the weighted harmonic mean area of the unguitractor apodeme (UTA) varies with the free length, too, the exact relationship between spring constant and length is complex, *k* = *EA*(*L*)*L*^−1^, and is evaluated here using data on the variation of *A* with *L* alongside the elastic modulus measurements for each tested apodeme (Figure 4 c and 5 b). The solid black line indicates a representative variation, extracted from the raw data via an arc-length analysis (Hartlen and Cronin, 2022); the dashed line indicates the variation expected for a sample with equal modulus but constant cross-sectional area, equal to the arithmetic average of the tested apodemes for comparison. Grey bars on the top indicate the average length of major limb segments for a hypothetical UTA with a length of 42 mm, corresponding to the UTA modelled by the arc-length analysis. n=13; colours indicate sample ID.

The variation in spring constant across the UTA length, though partially a direct mechanical consequence of the decrease in free length, may also be functionally interpreted with respect to the muscles that pull on it. The femoral retractor muscle actively engages the claws, and a lower apodeme spring constant may aid elastic recoil in the absence of an antagonist (Ichikawa et al., 2016; Radnikow and Bässler, 1991); the depressor and levator muscles in the tibia, in turn, move the proximal tarsomeres (Ichikawa et al., 2016), and a typically very small muscle in the distal part of the tibia is thought to be responsible for fine control (Radnikow and Bässler, 1991), functionally consistent with a larger spring constant and thus more immediate response.

## Discussion

In both invertebrates and vertebrates, muscle action is often mediated by elastic elements that connect muscle with the skeletal elements it actuates. Although these elastic elements are passive in the sense that they can generate neither force nor do work, they are thought to confer remarkable functional benefits to the mechanical performance of vertebrate musculoskeletal systems; their mechanical properties, crucial determinants of these benefits, have consequently been subject of a large body of work (e. g. Bennett et al., 1986; Bennett and Stafford, 1988; Ker, 1981; Lieber et al., 1991; Mossor et al., 2020; Pollock and Shadwick, 1994a,b; Shadwick, 1990; Shadwick et al., 2002; Summers and Koob, 2002). In sharp contrast, much less is known about the mechanical properties of the invertebrate tendon analogue—apodemes. Do they bring the same benefits? To increase our understanding of the function of in-series elastic elements in the most diverse animal group, we measured the modulus, ultimate tensile strength, and spring constant of the unguitractor apodeme (UTA) of sunny stick insects. Measuring these mechanical properties is a problem in metrology, the solution of which provides little biological insight by itself. Such insight must instead come from a comparative and functional analysis—a task we now tackle in the discussion.

### Apodemes and tendons have substantially different material properties

Sunny stick insects UTAs have a modulus of 8.03 ± 1.27 GPa, and a maximum tensile strength of 435 ± 158 MPa, both comparable to previous measurements for a different species, conducted by Bennet-Clark (E=18.9 GPa, *σ*_max_ ≈ 620 MPa Bennet-Clark, 1975), and Ker (E=8.1 and 13.2 GPa, respectively Ker, 1977). The modulus of insect limb apodemes thus appears to be of order 10 GPa; their tensile strength is roughly a factor 20-30 lower. To put these values into perspective, we collated modulus data from 22 vertebrate species of various body sizes (Dimery et al., 1986; Foutz et al., 2007; Jansen et al., 1998; Ker et al., 1988; Matson et al., 2012; Pollock and Shadwick, 1994b; Shadwick, 1990), focussing on the plantaris, the deep digital flexor, and the gastrocnemius tendons—major limb tendons thought to convey functional benefits during cyclical locomotion in cursorial and ambulatory vertebrates. These vertebrate tendons have an average modulus of 1.3 GPa (95 % CI [1.2; 1.4]), independent of an animal’s body mass (ordinary least squares regression on log10-transformed data, *t* = −0.67, *p* = 0.51, Figure 7 a)—about one order of magnitude smaller than the best estimate for the apodeme modulus (Figure 7 a).

**Figure 7.**
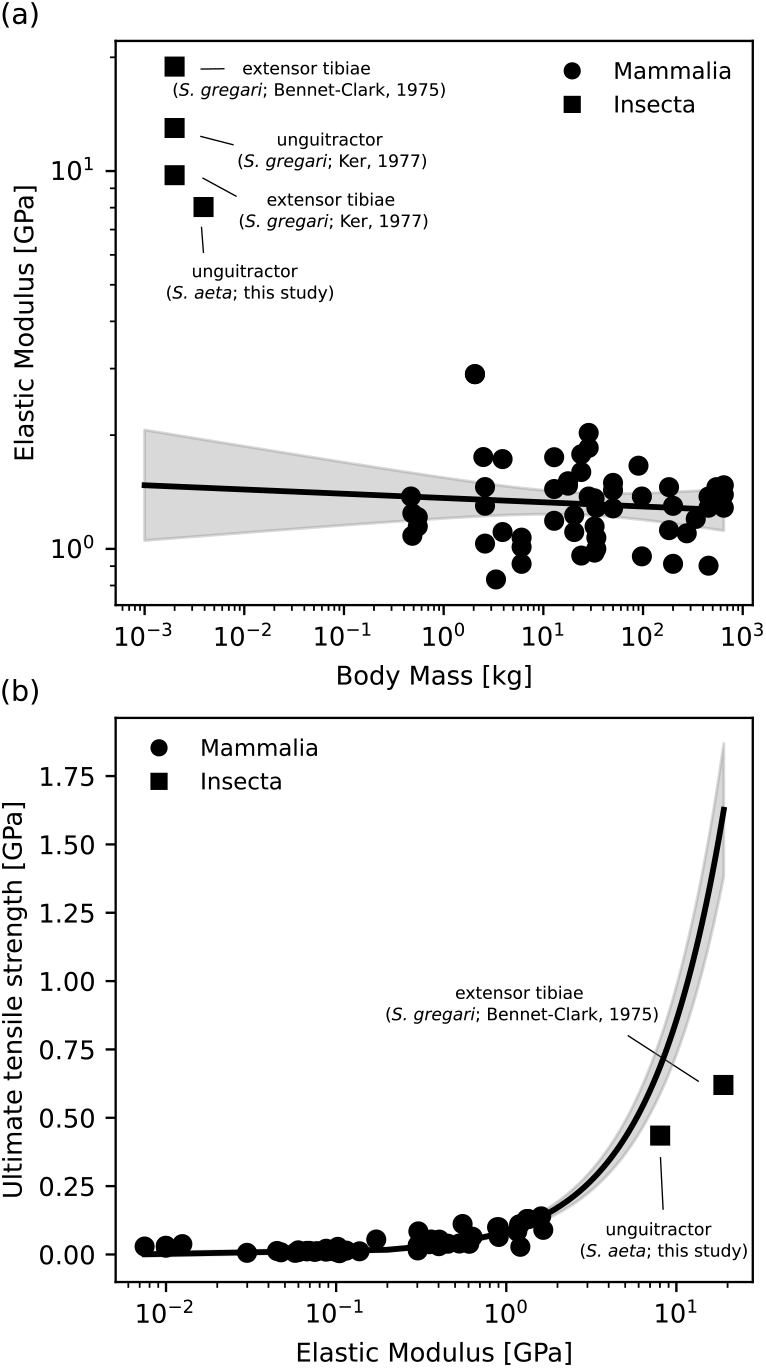
(a) The elastic modulus of insect apodemes— measured for *Sungaya aeta* stick insects and *Schistocerca gregari* locusts—appears to be about one order of higher than that of plantaris, deep digital flexor, and gastrocnemius tendons (size-invariant modulus of 1.3 GPa. Data from Bennet-Clark, 1975; Dimery et al., 1986; Foutz et al., 2007; Jansen et al., 1998; Ker, 1977; Ker et al., 1988; Matson et al., 2012; Pollock and Shadwick, 1994b; Shadwick, 1990). Note that both cuticle and collagenous tendons can be much softer and much stiffer (Abourachid et al., 2023; Labonte et al., 2017; Mossor et al., 2020; Politi et al., 2019; Stamm et al., 2021; Vincent and Wegst, 2004), suggesting that this difference is not merely a consequence of the material toolkit available to invertebrates and vertebrates. (b) Modulus and ultimate tensile strength (UTS) are usually correlated. To evaluate the UTS of insect apodemes relative to their modulus, we estimated the relationship between modulus and UTS for a wide range of in-series elastic tissues in vertebrates (data from Mossor et al., 2020), extrapolate this relation to the modulus of insect apodemes, and conclude that they seem to be weaker relative to their modulus. In both (a) and (b), solid lines show the result of ordinary least squares regressions on Mammalian data only, alongside the 95% confidence intervals (shaded); the regression in (b) was forced through the origin. Note the double-logarithmic axes in (a), and the semi-logarithmic plot in (b) (regression for (b) conducted on untransformed values).

The difference in modulus complicates the comparative analysis of the ultimate tensile strength, for modulus and strength typically correlate (Ashby and Jones, 2012). So, are apodemes unusually strong, weak, or exactly as strong as expected, given their modulus? To answer this question, we used the data assembled by Mossor et al. on the modulus and strength of vertebrate in-series elastic elements (Mossor et al., 2020), including not only various tendons, but also suspensory ligaments to characterise the relationship across a maximum range of moduli. The relationship between the modulus and strength of vertebrate elastic elements can be satisfactorily described via *σ*_max_ = 0.086*E* (ordinary least squares regression with zero intercept on untransformed data, 95 % CI: [0.076; 0.095]; R^2^ = 0.85; Fig. 7 b), close to if slightly higher than the classic first order prediction *σ*_max_ = 0.067*E* (Ashby and Jones, 2012). Extrapolated to a hypothetical vertebrate tendon with modulus of 8 GPa, the expected ultimate tensile strength is thus 687 MPa (95 % CI: [610; 765] MPa), significantly higher than the ultimate strength of stick insect UTAs (435 MPa ; 95 % CI: [339; 530] MPa. *t* = −438, *p <* 0.001). The disparity is more extreme still for locust apodemes, where extrapolation yields an estimate almost three times larger than the experimentally determined value (1618 vs 620 MPa, respectively). Thus, if anything, insect apodemes appear to be relatively weaker than their vertebrate analogues (Figure 7 b).

The marked difference in modulus and strength between tendons and apodemes is noteworthy, for material properties are usually assumed size- and taxon-invariant in functionally equivalent tissues, an assumption sometimes called “isophysiology” (Labonte and Holt, 2024; Polet and Labonte, 2024); as one illustrative example, the maximum isometric muscle stress does not deviate much from a typical value of about 350 kPa, at least within vertebrates (Marden, 2005; Marden and Allen, 2002; Medler, 2002). Some of the difference could be owed to measurement inaccuracy: to the best of our knowledge, all previous work on vertebrate tendons estimated the modulus as the ratio between the arithmetic mean stress and strain, and the strength as the ratio between force at failure and the arithmetic mean area (the most widespread method to determine the tendon cross-sectional area is to weigh the tendon, assume a density to calculate tendon volume, and then divide this volume by the measured tendon length, yielding the arithmetic mean area, see Bennett et al., 1986; Dimery et al., 1986; Javidi et al., 2019; Ker, 1981; Matson et al., 2012; Pollock and Shadwick, 1994a,b; Shadwick, 1990); where the tendon crosssectional area varies along the tendon length, this choice will likely result in a systematic underestimation of both modulus and strength. In the same vein, we note that apodeme ultimate strength measurements have remained challenging precisely because apodemes rapidly turn brittle *ex vivo*; it is consequently more likely that experiments result in an under- than in an overestimate of their true tensile strength (Bennet-Clark, 1975; Brown, 1963; Ker, 1977). However, it appears improbable that this experimental difficulty and a systematic error due to area variations would account for a difference of almost an order of magnitude, and the strength-to-modulus ratio for stick insect UTAs, *E/σ*_max_ = 18, remains very close to the first order prediction, *E/σ*_max_ = 15 (Ashby and Jones, 2012), lending some credence to its accuracy. As an alternative explanation, one may reasonably attribute the difference in material properties to the difference in material make-up: technically ingrowths of the exoskeleton, apodemes are derived from the same material tool kit, including chitin fibres and cuticular proteins; tendons, in turn are made from collagen and non-collagenous proteins. While the modulus-to-strength ratio may indeed simply differ between collagenous tendons and cuticular apodemes, it is arguably implausible that the absolute difference in modulus itself is a result of taxon-specific material limitations: cuticle is a remarkably diverse material, and can have moduli orders of magnitude lower, and at least a factor of two higher (Labonte et al., 2017; Politi et al., 2019; Stamm et al., 2021; Vincent and Wegst, 2004). The tendon modulus, too, can be much higher or much lower than the reported average, provided specialisation calls for it, for example in the calcified tendons of some birds (modulus up to 16 GPa Bennett and Stafford, 1988), where a high modulus aids tendon function during a posture stabilisation based on tensegrity (Abourachid et al., 2023), or in the tendons of sloths, which are so compliant that they appear more akin to suspensory ligaments (modulus as small as 0.07 GPa, Mossor et al., 2020). The most plausible interpretation of the observed difference in moduli, then, is that it reflects a genuine difference in selected material properties. What are the implications of this difference for the functional significance of in-series elastic elements in vertebrate versus invertebrate musculoskeletal systems?

### Comparative allometry of the spring constant of in-series elastic elements

Virtually all supposed functional benefits of in-series elastic elements demand that they stretch by a Goldilocke amount—too compliant or too stiff are ill-suited both. Colloquially, both the spring constant, *k*—the proportionality constant between force and displacement, *k* = *F/δ*— and the modulus, *E*—the proportionality constant between stress and strain, *E* = *σ/ε*—are used as indicators of stiffness. However, *k* and *E* differ crucially in two aspects: First, while a high spring constant always implies high stiffness by definition, a material with a high modulus can have high or miniscule stiffness, for the stiffness depends on both modulus and specimen geometry, *k* = *EAL*^−1^. In other words, a short and thick specimen with low modulus and a thin and long specimen with a high modulus may well have the same spring constant. Second, and as a direct consequence, the spring constant retains a size-dependence, *k* = *EAL*^−1^ ∝ *m*^1*/*3^ in isogeometry and isophysiology. Thus, even if invertebrate apodemes were exact scaled-down versions of vertebrate tendons with identical modulus, they would nevertheless be substantially less stiff in absolute terms. To avoid errors in the comparative and functional analysis of in-series elastic elements, it is thus crucial to maintain a clear distinction between the material and the functional system property, i. e. the modulus and the spring constant, respectively.

One statistical approach to compare apodeme and tendon spring constants proceeds through assessment of the allometry of tendon spring constants. With this relation at hand, the spring constant of a hypothetical vertebrate with the body mass of a sunny stick insect may be inferred from extrapolation and compared to the measured value. To perform such an analysis, we used data on the arithmetic mean cross-sectional area, total length and modulus of the plantaris, the deep digital flexor, and the gastrocnemius tendons of 18 quadrupedal vertebrates ranging from 470 g to 545 kg in body mass (Pollock and Shadwick, 1994a,b, where the modulus was missing in the original data, we used the arithmetic mean of *E* = 1.3 GPa, derived from the meta-analsysis above). Estimating the *in vivo* tendon spring constant then requires definition of the tendon’s free length. In the vertebrate literature, this problem is often dealt with by invoking an “effective tendon length”, equal to the difference between muscle fibre and tendon length (Alexander and Vernon, 1975; Ker et al., 1988; Pollock and Shadwick, 1994a). For the tendons considered here, the fibre length is so small that the effective length differs little from the total tendon length (Pollock and Shadwick, 1994b), so that, in service of simplicity, we neglected it altogether, *k* = *EAL*^−1^. An ordinary least-squares regression on log10-transformed data then yields *k* = 26828 N m^−1^ m^0.3^ (with mass in kilograms, 95% CI of scaling coefficient [0.23; 0.36]), consistent with isogeometry, *k* ∝ *m*^1*/*3^ (Figure 8 a). Extrapolating this allometric relation to a vertebrate with a body mass of 3.9 g, equal to the mean body mass of our stick insect specimens, yields an estimated spring constant of 5206 N m^−1^, (95% CI [2856; 9490] N m^−1^; Figure 8 a)—a value more than 10 times larger than the minimal UTA spring constant calculated from the weighted harmonic mean cross-sectional area and the total apodeme length (319 N m^−1^ (95% CI: [272; 366] N m^−1^)). However, the effective free length of the UTA is likely much smaller than its total length, as muscles attach all along the proximate half of the femur, and at several positions along the length of the tibia. Reevaluating the weighted harmonic mean area for these two free lengths, defined as the distance between the unguitractor plate attachment and the distal end point of muscle attachments in femur and tibia, respectively, yields spring constants of 400 and 7000 N m^−1^ (Figures 8 a-c))— spanning the extrapolated estimate from the mammalian tendon regression. It would appear that the spring constant of UTAs is about as high as expected given the size of the animals; if anything, it is somewhat lower.

**Figure 8.**
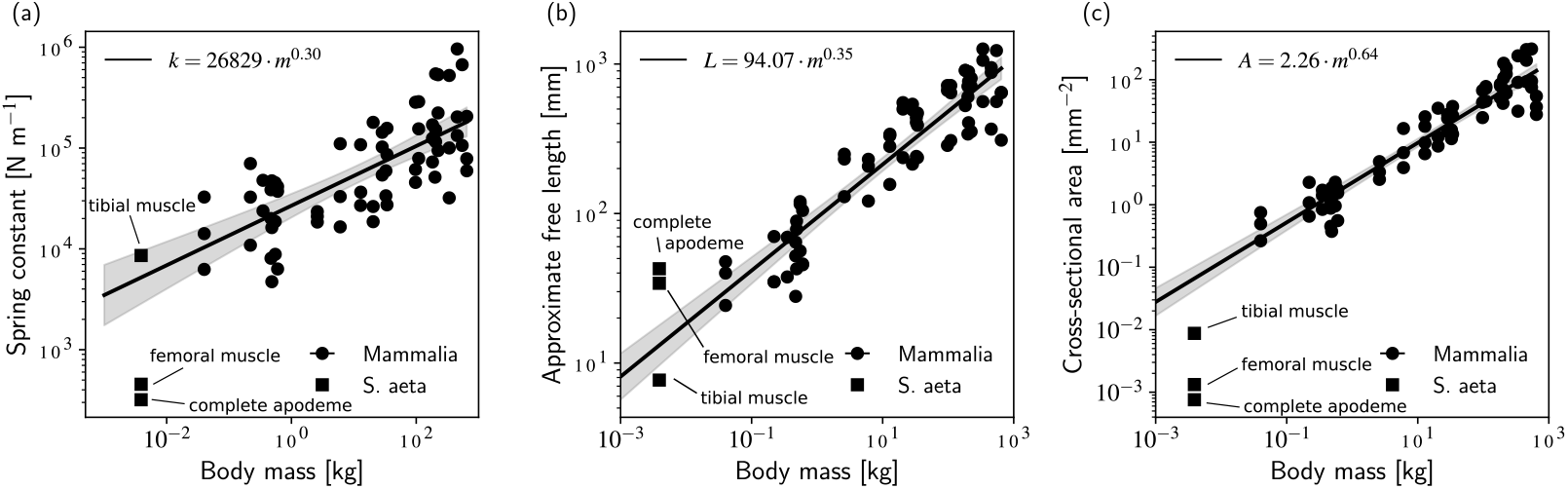
(a) The spring constant is a system parameter, which depends on both material properties and geometry, *k* = *EAL*^−1^. One consequence of this dependency is that a specimen with equal geometry and material properties, downscaled isometrically, will have a smaller spring constant than its larger twin, *k* ∝ *m*^1*/*3^. Assessing the functional significance of the unguitractor apodeme thus requires a correction of the measured spring constant for size. To conduct such a correction, we extrapolated the relationship between tendon spring constant and body mass in mammals, extracted from Pollock and Shadwick (1994b), to the body mass of a *Sungaya aeta* stick insect; it may then be compared to three spring constant estimates: (i) the spring constant of the complete apodeme, against which external forces pull; (ii) the effective spring constant experienced for muscles in the femur; or (iii) the tibia. Remarkably, although the UTA modulus is about an order of magnitude larger than the average modulus of major limb tendons (Figure 7 a), its absolute and size-corrected spring constant is, if anything, lower; it approaches the extrapolation only for the short effective length experienced by muscles in the tibia. This difference arises because (b) length and (c) cross-sectional area depart from the prediction via extrapolation, too: UTAs appear to be longer and thinner than expected from isometric extrapolation, which partially compensates for the difference in modulus. This many-to-one-mapping of morphology and material properties to spring constant suggests that apodemes may have a normalised spring constant comparable to that of tendons, and may thus bring the same functional benefits (for a more detailed discussion, see text). The solid lines are the results of ordinary least square regressions on log10-transformed data for mammalia only; the free length estimates for *S. aeta* are approximate, assuming muscles attach in the proximal half of the femur, and across the entire tibia, as estimated via light microscopy.

### The functional relevance of apodemes, and many-to-one mapping in musculoskeletal design

A size-specific apodeme spring constant that is, if anything, *lower* than that of vertebrate tendons may come as a surprise to those familiar with the literature. A considerable body of work has asserted that apodemes are so rigid that one may (i) assume instantaneous force transfer (e. g. Miles et al., 1995); (ii) consequently infer muscle strains directly from joint angles (e. g. Ahn and Full, 2002; Ahn et al., 2006; Full et al., 1998); and (iii) exclude them from theoretical work altogether (e. g. Kukillaya and Holmes, 2009; Schmitt and Holmes, 2003; Zakotnik et al., 2006). How did this idea take hold?

The notion that invertebrate apodemes are much stiffer than vertebrate tendons can eventually be traced to a pioneering study by Full and Ahn (Full and Ahn, 1995). In what was probably the first numerical musculoskeletal model of an insect limb, the authors estimated that insect apodemes are about 40 times stiffer than vertebrate tendons (Full and Ahn, 1995, eq. 6; p. 1290; note that the authors pointed out that this estimate is likely too high.). In search of the origin of the difference between this and our assessment, we first note that Full and Ahn refer to a *normalised* apodeme spring constant: following Zajac (1989), they divided *k* with a characteristic muscle force, *F*_*m*_, and multiplied the result with the optimal muscle fibre length, *L*_*m*,0_, to find a non-dimensional spring constant, 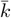:

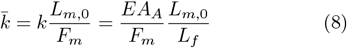

where *A*_*A*_ is a characteristic apodeme cross-section, and *L*_*f*_ is the unstretched (slack) length of the apodeme. Using Ker’s value of *E* = 13 GPa (Ker, 1977), and an estimate for the apodeme stress of *F*_*m*_*/A*_*A*_ ≈ 8.6 MPa, Full and Ahn arrived at 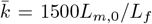, which they compared to an estimate for the average human limb tendon, 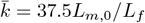, from Hoy et al. (1990); from here, one may find 1500*/*37.5 = 40—the origin of the factor 40. However, note well that this comparison tacitly assumed that the normalised tendon and apodeme lengths are equal (the term *L*_*f*_ */L*_*m*,0_ in eq. 8). To put this assertion to the test, we again draw on Pollock and Shadwick (1994a) to estimate a median normalised tendon length of about 13; from table 2 in Full and Ahn (1995), we in turn extracted a median normalised apodeme length of 0.31—a difference of almost exactly a factor 1/40. This naive estimate thus suggests that the normalised tendon and apodeme lengths may differ by just the factor that is needed to balance out the difference in *EA*_*A*_*/F*_*m*_— apodemes and tendons may well have the same normalised spring constant.

Four sources of uncertainty make it prudent to treat this conclusion with great caution. First, the estimate of the apodeme cross-sectional area in Full and Ahn (1995) cannot be reconstructed with certainty: *A*_*A*_ is assumed to be proportional to the muscle physiological cross-section, but, to the best of our knowledge, the proportionality constant itself remains unspecified (Full and Ahn, 1995, p. 1290). Second, the estimates for *EA*_*A*_*/F*_*m*_ ≈ *E/σ*_*A*_ are based on one cockroach species and a selection of human limb tendons, and are thus of limited robustness— an issue that of course also afflicts our data, which draws from but two insect species, and which focuses on what is likely the longest, thinnest and thus least stiff invertebrate apodeme. Third, after recognising that the ratio *E/σ*_*A*_ is, in fact, the inverse of a characteristic apodeme strain, one may reasonably wonder if the substantial difference in strain across vertebrates and invertebrates that these estimates imply—and an absolute apodeme strain as low as 100*/*1500 = 0.067%—are all that plausible. Why would selection have ignored a design so inefficient as to operate with safety factors as high as 435*/*8.6 ≈ 50, at least a factor 10 larger than estimates across a large range of biological materials (Alexander, 1981)? Fourth, and as perhaps the most salient point, the normalisation suggested by Zajac (1989) may follow straight from dimensional analysis, but we argue that it is functionally incomplete, precisely because it omits a characteristic strain—that of the muscle that pulls on the apodeme. To justify this claim, we point out that the two spring constants implicit in eq. 8 are defined differently: the apodeme spring constant reads *k*_*A*_ = *EAL*^−1^, but the effective muscle spring constant is introduced as *k*_*M*_ = *σAL*^−1^—one relies on a modulus, the other on a stress. Their ratio thus returns a strain, 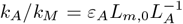, which should be accounted for by division with the muscle strain, *ε*_*M*_, to derive a functionally consistent definition of 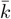 (the same conclusion may be derived at via an energy balance). Even this result is not unique, however, for dynamic contractions may well call for a different normalisation altogether, involving the payload mass and muscle strain rate instead (Galantis and Woledge, 2003)—further theoretical work is required to identify the functionally relevant normalisation for the spring constant of in-series elastic elements.

Regardless of the appropriate form of eq. 8, one conclusion appears clear enough: the same normalised spring constant can be achieved through substantially different means. For example, although stick insect apodemes may have a much higher modulus than vertebrate tendons, their size-specific spring constant is nevertheless comparable, because the difference in modulus is counteracted by a roughly equivalent difference in apodeme geometry. Indeed, whatever the form of eq. 8, it will ultimately encode a many-to-one mapping (e. g. Alfaro et al., 2005; Arnold, 1983; Blanke et al., 2018; Chatar et al., 2022; Moen, 2019; Polet and Labonte, 2024; Strobbe et al., 2009): non-trivial variations in musculoskeletal anatomy and physiology can achieve equivalent relative spring constant across very different “designs”. The noteworthy evolutionary consequence of such many-to-one mapping is the possibility of morphological and physiological diversification despite a convergence in mechanical function (Muñoz, 2019; Muñoz et al., 2018; Wainwright, 2007; Wainwright et al., 2005); viewed in this light, the difference in modulus across vertebrates and invertebrates could well be a mere phylogenetic accident.

In light of our results, the assertion that insect apodemes are functionally stiffer than their vertebrate analogues—perhaps even so stiff that they contribute to musculoskeletal performance in fundamentally different ways—must be considered pre-mature. Why, any such claim ought to be subjected to an extraordinarily high burden of proof, for it remains unclear what mechanisms, evolutionary or physically, would have prevented the largest and most diverse animal group on this planet from exploiting the considerable benefits ascribed to stretchable in-series elastic elements in vertebrates. A thorough assessment of apodeme function requires rich comparative work that integrates material science (modulus, strength), morphology (cross-sectional areas, lengths), physiology (muscle stress, strain and strain rate), and theoretical modelling (what is the appropriate normalisation?); such work will ultimately increase our understanding of the comparative biomechanics of musculoskeletal systems.

## Acknowledgments

The authors thank Prof. Dr.-Ing. Jörg Müssig for his insightful advice on tensile testing challenging specimen, and all members of the Evolutionary Biomechanics Laboratory at Imperial for their support and kindness.

